# How does co-occurrence of *Daphnia* species affect their gut microbiome?

**DOI:** 10.1101/2024.09.10.612237

**Authors:** Shira Houwenhuyse, Francois Massol, Emilie Macke, Luc De Meester, Isabel Vanoverberghe, Robby Stoks, Ellen Decaestecker

**Affiliations:** Ghent University and KULAK; CILL; KULAK; IGB; KULeuven

**Keywords:** *Daphnia*, microbiome, bacterioplankton, co-occurrence, competition

## Abstract

Species co-occurrence can lead to competitive interactions that influence fitness. Competition is typically assumed to be modulated by species niche, especially food–acquisition related traits. The influence of interspecific interactions on host microbiome communities has rarely been considered, and yet may provide an alternative mechanism regarding the effect of host species co-occurrence on their fitness. Here, we investigated whether the composition of the gut microbial community differs between two *Daphnia* species (*D. magna* and *D. pulex*), and whether the gut microbiome of one species depends on the presence of the other. We hypothesized the stronger filter-feeder *D. magna* to have a larger effect on the gut microbiome of the weaker filter-feeder *D. pulex* than *vice versa*. To this purpose, three *D. magna* and three *D. pulex* genotypes were first made axenic and then grown in monocultures or in cocultures in natural environmental bacterioplankton-enriched water, before assessing the community composition of the gut microbiomes and bacterioplankton. We found that the composition of the gut microbiome of the two *Daphnia* species did not significantly differ overall. However, subtle differences between mono- and cocultures were found at the *Daphnia* genotype level. For most genotype combinations (six out of nine), the microbiome of *D. pulex* changed more when grown in cocultures with *D. magna* than in monocultures. This provides limited support for our hypothesis that the stronger competitor has a larger effect on the gut microbiome of the weaker one than *vice versa*, and that this effect is possibly mediated via the bacterioplankton community.

## Introduction

Interspecific competitive interactions can play an important structuring role in communities via their effects on organismal growth and survival. An often-overlooked factor regarding how such interactions may shape life-history traits is through the microbiota associated with the competing host species (Augustin et al., 2017). Microbiota can contribute to host fitness by affecting host digestion, metabolism, growth and immunity (Franzenburg et al., 2013; Motiei et al., 2020). Microbiomes can, also contribute to host competitiveness. Host-associated bacteria can alter the strength of host interspecific interactions by facilitating host growth in the presence of an established species (Jackrel et al., 2020). Yet, *vice versa*, competition among host species may also shape microbiomes, for example by changing the environmental pool of bacteria (Razak et al., 2019; Macke et al., 2020). Host-associated factors may influence gut microbiomes, but conversely the microbiome may also affect host performance and possibly its environment via direct and indirect effects (Berendson et al., 2012; Macke et al., 2020; Weiss and Aksoy, 2011; Decaestecker, Van de Moortel et al., 2024). Disentangling the factors affecting the composition of host microbiomes is an unresolved puzzle at the interface between genetic and environmental processes. From the diverse pool of environmental microbes, specific micro-organisms are recruited in the host depending on the host immunity and genetic background, as well as on complex interactions between microbes, host physiology and particular environmental conditions (Goodrich et al., 2004; Engel & Moran, 2013; Chassaing and Cascales, 2018; Zélé et al. 2018; Halliday et al. 2020; Macke et al. 2020), including the presence of competing host species (He et al., 2018).

Species-specific microbiomes can influence the gut microbiome of co-occurring species, for example when microbiota are exchanged via the environment (Avena et al., 2016; Sehnal et al., 2021). There is evidence both in plants (Berendson et al., 2012) and in animals (e.g., Reveillaud et al., 2014; Engel et al., 2018; Fraune and Bosch, 2007) for species-specific microbiomes.

Interspecific variation in the niche, social behavior, and geographic range of the host can generate differences in the gut microbiota composition among species by altering the pool of microbial species that the host encounters and the patterns of microbial transmission (Adiar et al., 2020). On the other hand, even when co-occurring species have a similar microbiome, an influence on the other host’s microbiome is possible, for example if one species changes the bacterial species pool that can be taken up by the other species. This can be expected when one species has a higher grazing rate on bacteria, leaving more grazer-resistant strains in the environment, which in turn can be taken up by the other species.

Rarely has the influence of interspecific interactions on host microbiome communities been considered. He et al. (2018) explored the effect of interspecific competition on the gut microbiome of two Atlantic salmon populations. They found that interspecific interactions can cause a lower alpha diversity of gut microbiota, loss of beneficial bacteria, and a larger change in microbial community in the weaker competitor of two Atlantic salmon species. Effects on the gut microbiome of the weaker competitor were probably related to competition-related stress, resulting in the decrease or loss of beneficial bacteria. Competing host species can also exchange microbiota. This can in turn induce interspecific microbiota-mediated facilitation effects (Zélé et al., 2018). Facilitative priority effects occur when a species (e.g., a bacterium) that arrives first at a site alters the (a)biotic conditions in ways that positively affect a later arriving species (Halliday et al., 2020; Decaestecker, Van de Moortel et al., 2024).

Investigation of the effects between competing host individuals and their microbiomes requires host species (i) that are relatively phylogenetically close, ideally congenerics, as these are more likely to share microbiota, (ii) with an easy-to-manipulate microbiome so that axenic organisms can be obtained, and (iii) with contrasted competitive abilities. The freshwater crustacean genus *Daphnia* fits these criteria nicely and has been successfully used to study the interplay between *Daphnia* species, their microbiomes and the environment. Several bacterial taxa are consistently found in affiliation with *Daphnia* (Proteobacteria, Actinobacteria, and Bacteroidetes; Motiei et al., 2020), even in geographically separated cultures (Qi et al., 2009). Microbiomes of *D. magna* genotypes from different regions are distinct, even years after being brought into the laboratory (Frankel-Bricker et al., 2019; Houwenhuyse et al., 2021). Many recent studies show that the *D. magna* bacterial community is structured differently than that from the surrounding bacterioplankton and from the microbiome of their food (Freese and Schink, 2011; Berg et al., 1016; Callens et al., 2016; Cooper and Cressler; 2020; Motiei et al., 2020). The gut microbiome of *D. magna* also changes with diet, suggesting a strong influence of the environment on the bacterial community in the gut (Freese and Schink, 2011; Callens et al., 2016; Macke et al., 2017, Eckert et al., 2021), and thus the presence of a flexible microbiome. The difference between the microbiome found in the gut of *D. magna* and in its environment suggests that the host and its associated microbes interact to establish and maintain these microbial populations.

In freshwater ecosystems, the competition between different-sized *Daphnia* species can be intense because of their similar life strategies and overlap in feeding habits (Chen et al., 2016). *Daphnia magna* and *D. pulex* are two frequently co-occurring cladoceran species in freshwater ponds, reservoirs and lakes (Kuster and Von Elert, 2013; Asselman et al., 2014). Compared to *D. pulex*, *D. magna* is larger-bodied, a stronger filter-feeder, faster-growing, and more competitive, but also more vulnerable for microparasite exposure (Decaestecker et al., 2005; 2015). According to the size-efficiency hypothesis (Brooks and Dodson, 1965), larger-bodied zooplankton species are competitively superior because they are more efficient filter-feeders, they can ingest more and larger food particles, including bacteria and algae, and they have relatively reduced metabolic demands per unit mass than smaller-bodied species.

The aim of this study is to determine whether the composition of the gut microbial community differs between *D. magna* and *D. pulex*, and whether the microbiome depends on the presence of the other species. We hypothesized the stronger filter-feeder *D. magna*, by changing the environmental bacterioplankton pool, to have a larger effect on the gut microbiome of the weaker filter-feeder *D. pulex* than *vice versa* (Figure 1). To this purpose, multiple maternal lines of three *D. magna* and three *D. pulex* genotypes were made germ-free, and then placed in water enriched with natural environmental bacterioplankton. Each genotype was grown in mono- and in coculture and the community composition and diversity of the gut microbiomes and bacterioplankton were investigated. We assessed (i) differences in the gut microbiome composition between the *Daphnia* species, (ii) the effect of the presence of another host species on gut microbiomes by comparing microbiome composition in mono- versus coculture, and (iii) the asymmetry of these effects, testing whether *D. magna* had a stronger effect on *D. pulex* than *vice versa*.

**Figure 1:**
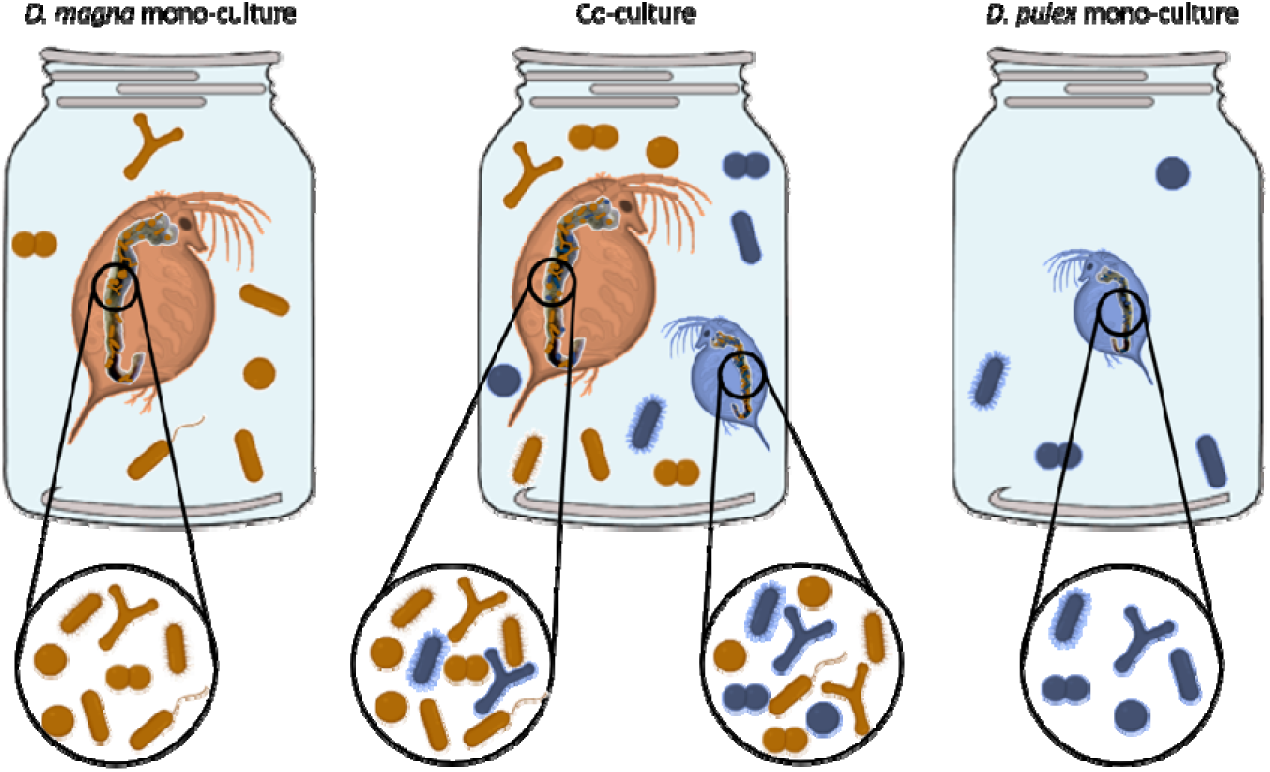
Graphical presentation of our hypotheses that (i) the gut microbiome composition differs between *D. magna* (orange) and *D. pulex* (blue), (ii) the gut microbiome differs in monocultures versus cocultures, and (iii) the larger filter-feeder *D. magna* has a stronger effect on the gut microbiome of the smaller filter-feeder (*D. pulex*) than *vice versa*.

## Materials and methods

### *Daphnia* and algae culturing

To investigate whether the composition of the gut microbial community depends on the presence of the other host species, three *D. magna* genotypes (KNO15.04, OM2NF4 and T2) and three *D. pulex* genotypes (DB, DK, GB) were cultured in the laboratory (Table 1). Each genotype came from a different natural population in Belgium where *D. magna* and *D. pulex* co-occur. All genotypes were maintained in the laboratory under standardized conditions for several years prior to the experiment. Stock *Daphnia* clonal lineages were cultured in filtered tap water at a temperature of 19 ± 1°C and under a 16:8h light:dark cycle in 2L glass jars (at a density of 30 individuals/L). They were fed three times/week with saturating amounts of the green algae *Chlorella vulgaris*. Four iso-female lines were established as four independent maternal lines per clone in preparation for this experiment. These iso-female lines were fed every day and medium was refreshed once per week.

**Table 1:** Geographic locations of the genotypes used in this experiment.

| Genotype | Species | Location | Coordinates |
| --- | --- | --- | --- |
| KNO15.04 | *D. magna* | Knokke | 51°20'05.62"N, 03°20'53.63"E |
| OM2NF4 | *D. magna* | Heverlee, Abdij van 't Park | 50°51'45.0"N, 04°42'58.8"E |
| T2 | *D. magna* | Oud-Heverlee | 50°50'24.0"N, 04°39'40.4"E |
| DB | *D. pulex* | Destelbergen | 51°02'49.4"N 3°48'33.3"E |
| DK | *D. pulex* | Donkmeer | 51°02'18.5"N 3°58'37.6"E |
| GB | *D. pulex* | Geraardsbergen | 50°46'17.6"N 3°52'56.6"E |

*Daphnia* were fed with the unicellular green algae *C. vulgaris* during the experiment. The green algae *C. vulgaris* is good-quality food for *D. magna* (Munirasu et al., 2016). *Chlorella vulgaris* was grown in Wright’s Cryptophyte medium under sterile conditions in a climate chamber at 22 ± 1°C with a light:dark cycle of 16:8h in 2L glass bottles, with constant stirring and aeration. Filters (0.22 µm) were placed at the input and output of the aeration system to avoid any bacterial contamination. The algae were harvested weekly in the stationary phase.

### Experimental design

For each genotype, four iso-female lines were grown in separate jars for two generations to control for maternal effects. Growth conditions of these four iso-female lines per genotype were as in the stock cultures. Eggs from the second brood were collected and disinfected with peracetic acid, following the protocol of Callens et al. (2016). Thirty axenic juveniles were then collected and divided into two jars: (1) monocultures of the genotype (20 individuals/jar) and (2) cocultures with a genotype from the other species ((10 *D. magna* + 10 *D. pulex*)/jar; Figure 2). The total number of *Daphnia* individuals/jar was standardized over the different treatments.

**Figure 2:**
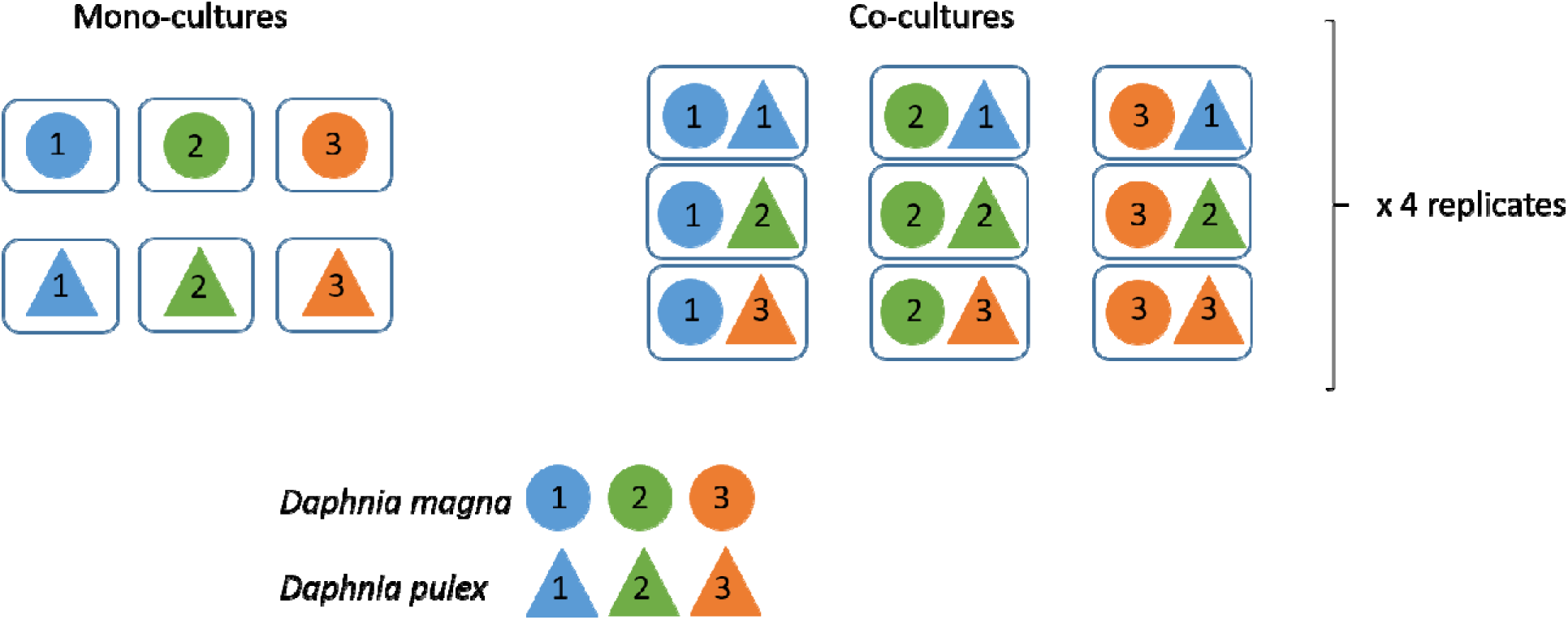
Experimental design to study the effect of being cultured in mono-versus cocultures on the gut microbiome of *D. magna* (represented by circles) and *D. pulex* (represented by triangles). Three genotypes (numbers within symbols) were studied per species (see Table 1). These genotypes were kept in mono- and cocultures for 10 days. In both, mono- and cocultures the total number of *Daphnia* individuals was 20, i.e., in monocultures 20 individuals *D. magna* or *D. pulex*, in cocultures 10 *D. magna* + 10 *D. pulex* individuals. For each mono- and coculture, there were four replicates per combination of genotypes.

There were 15 treatments (six monocultures + nine cocultures, Figure 2) in total, with for each treatment four independent replicates, resulting in a total of 60 experimental jars. All experimental jars contained sterile filtered tap water and 20% of pond water (= pond water inoculum), collected from a natural pond in the Ecolab at the KU Leuven campus Kortrijk (50°48’18.8”N, 3°17’34.3”E). The pond water inoculum of each replicate was sequenced to investigate the bacterial community present. *Daphnia* individuals were maintained under these conditions for ten days. They were then placed in sterile filtered tap water for 12h to remove food particles from the gut, as well as environmental bacteria on the carapace and filter apparatus, after which their guts were dissected. For the monoculture treatment, 10 *Daphnia* individuals per jar were dissected and placed together in an Eppendorf tube. For the coculture treatment, 10 *D. magna* and 10 *D. pulex* individuals per jar were dissected, and their guts were placed in two separate Eppendorf tubes. In total, there were 96 samples containing guts (six monocultures x 4 replicate jars + nine cocultures x 2 species x 4 replicate jars). In addition, the bacterioplankton from each jar was collected by filtering approximately 100 mL over a 0.22 µm filter to examine the bacterioplankton community. There was a total of 60 bacterioplankton samples.

To obtain axenic recipient juveniles, the protocol described by Callens et al. (2016) was followed. Females carrying parthenogenetic eggs were dissected under a stereomicroscope with dissection needles. Per iso-female clonal line, a total of 90 eggs were collected by batches of 30 in a Petri-dish in 10 mL filtered tap water. Only recently deposited eggs, characterized by the presence of an external membrane, were isolated. The Petri-dishes with eggs were then placed in a laminar flow hood to disinfect the eggs by submersing them in 10 mL of a 0.01% peracetic acid solution and gently agitating them for 10 minutes. After 10 minutes in the disinfectant, the eggs were transferred to another Petri-dish, containing 10 mL sterile filtered tap water to remove peracetic acid residues. The eggs stayed in this rinsing step for 10 minutes after which the rinsing step was repeated, to ensure that any trace of peracetic acid was removed. Afterwards, the eggs were transferred in groups of 30 to wells of a 6-well plate filled with 8 mL sterile filtered tap water. The 6-well plate was sealed with parafilm and placed in an incubator. Eggs were allowed to hatch for 48 h under sterile conditions at a temperature of 20 ± 0.5°C and a 16:8h light:dark cycle.

### Library preparation and sequencing

To characterize the microbial communities of the *Daphnia* guts, DNA was extracted using a PowerSoil DNA isolation kit (Qiagen). DNA was dissolved in 20 µL milliQ water. The total DNA yield was determined using a Qubit dsDNA HS assay (Invitrogen) on 1 µL of sample. Because of initially low bacterial DNA concentrations in some samples, a nested PCR was applied to increase specificity and amplicon yield. The full-length 16S rRNA gene was first amplified with EUB8F and 1492R primers on 10 ng of template using a high-fidelity SuperFi polymerase (Life Technologies) for 30 cycles: 98°C – 10s; 50°C – 45s; 72°C – 30s. PCR products were subsequently purified using the QIAquick PCR purification kit (Qiagen). To obtain dual-index amplicons of the V4 region, a second amplification was performed on 5 µL (=20-50 ng) of PCR product using 515F and 806R primers for 30 cycles: 98°C – 10s; 50°C – 45s; 72°C – 30s. Both primers contained an Illumina adapter and an 8-nucleotide (nt) barcode at the 5’-end. For each sample, PCRs were performed in triplicate, pooled and gel-purified using the QIAquick gel extraction kit (Qiagen). An equimolar library was prepared by normalizing amplicon concentrations with a SequalPrep Normalization Plate (Applied Biosystems) and subsequent pooling. Amplicons were sequenced using a v2 PE500 kit with custom primers on the Illumina Miseq platform (KU Leuven Genomics Core), producing 2 x 250-nt paired-end reads.

### ASV sequence processing

DNA sequences were processed in R 4.0.0 (R studio version 1.1.463) following Callahan et al. (2016). Sequences were trimmed (the first 10 nucleotides and from position 190 onwards) and filtered (maximum of two expected errors per read) on paired ends. Sequence variants were inferred using the high-resolution DADA2 method, which relies on a parameterized model of substitution errors to distinguish sequencing errors from real biological variation. Chimeras were subsequently removed from the data set. After filtering, a total of 4,844,123 reads were obtained with on average 30,088 reads per sample (minimum = 0 read and maximum= 158,399 reads). To test for differences in α- and β- diversity, all samples were rarefied to a depth of 1,000 reads, based on the number of reads per sample and the rarefaction curve (Figure SI1). Sixteen samples had a lower number of reads and were removed from the analysis. Taxonomy was assigned with a naïve Bayesian classifier using the Silva v132 training set. ASVs with no taxonomic assignment at the phylum level or which were assigned as “chloroplast” or “cyanobacteria” were removed from the data set. ASVs for which the mean relative abundance was below 10^-5^ were also removed from the analysis. To visualize the bacterial families that differed between the treatments, ASVs were grouped at the family level, and families representing <1% of the reads were discarded. Close investigation of the four replicates showed that the gut microbiomes of replicate 4 deviated strongly from the others. Possibly the lower species richness in the start inoculum from replicate 4 (only 10 bacterial species compared to 26, 27 and 73 in respectively replicates 1, 2 and 3) caused the gut microbiomes of replicate 4 to group separately on the ordination plots. Because of this strong bias, we excluded replicate 4 from further analysis. The results with replicate 4 included can be found in the supplementary information (Figure SI1-SI6 and Table SI1-SI6).

### Statistical analyses of the microbiome

Measures of alpha diversity of the gut microbial communities within the different treatments, i.e. ASV richness and the Shannon diversity (taking into account both ASV richness and the relative abundance of ASVs) were calculated using the vegan package in R, following Borcard et al. (2011). Shannon diversity (Hill number of order 1) was calculated as the exponential function of the Shannon entropy, which represents the effective diversity of the bacterial community in the sample (Banos, 2006). Hill numbers (species richness, Shannon diversity and Simpson diversity; respectively Hill number of order 0, 1 and 2) were calculated following Hsieh et al. (2016).

Linear mixed-effects models (LMER) were used to model bacterial community diversity in terms of sample type (bacterioplankton vs. gut microbiota), culture type (mono- vs. coculture), species (*D. magna* vs. *D. pulex*), genotype, and the interactions of these factors, together with various factors affecting the structure of random effects (genotype nested in species and species nested in culture type). The Akaike information criterion (AIC) was used to select the best model (Table SI1). The importance of the fixed factors in the best model was tested with F-tests (Type II ANOVA) using Satterthwaite and Kenward-Roger methods for denominator degrees of freedom for *F*-tests and *p* values (Anova function of the car package). The best model of the full dataset (bacterioplankton and gut samples) included sample type (bacterioplankton versus gut sample), culture type (mono- versus coculture), and all possible interactions as fixed factors; genotype nested within species, and species nested within culture type were considered random factors. The best models (when separately analyzing the bacterioplankton and gut microbiomes) included culture type and all possible interactions as fixed factors with genotype nested within species, and species nested within culture type as random factors. Post-hoc analyses on coefficient estimators were performed using the ‘emmeans’ function with Tukey’s adjustment of p-values from the emmeans R package.

To investigate differences based on individual dissimilarities in community composition (beta diversity) among the microbial communities, Bray-Curtis dissimilarity indices, as well as weighted and unweighted Unifrac distances, were calculated and plotted using principal coordinate analysis with the phyloseq package in R. In addition, distances between bacterial communities harbored by the two different *Daphnia* species were plotted per genotype, comparing mono- versus cocultures. The effects of sample type (bacterioplankton or gut microbiomes), culture type (mono- or coculture), species (*D. magna* or *D. pulex*) and genotype, and all possible interactions, on dissimilarity were assessed through a non-parametric multivariate analysis of variance, using the Adonis2 function in the vegan package in R (Anderson, 2001).

Differential abundance analyses were performed to identify bacterial families that differed between groups (e.g., bacterioplankton vs gut microbiomes, mono- vs coculture, etc.) with the Bioconductor packages DESeq and EdgeR. This was done on the raw sequencing data (with removal of two samples that had 0 reads). With DESeq, pairwise comparisons were made, while with EdgeR interactions between the groups were investigated. The Benjamini-Hochberg adjustment was applied to correct for multiple testing in the DESeq and EdgeR analysis. To visualize the number of unique and shared ASVs between groups, Venn diagrams were created on the rarified data using the ggVenn package in R. Bacterioplankton and gut microbiomes were visualized separately, to investigate unique and shared ASVs between the two host species, *D. magna* and *D. pulex*.

All statistical analyses of the microbiome data were performed in R 4.0.0 (R studio version 1.1.463).

## Results

### Alpha diversity

Investigation of the full dataset (bacterioplankton and gut microbiomes) showed a significant difference in species richness (F=49.86; df=1, 67; p<0.0001) and in Shannon diversity (F=17.68; df=1, 67; p<0.0001) between gut microbiomes and bacterioplankton, with the bacterioplankton being in general more than three times less diverse than the gut microbiomes (Figure 3). The magnitude of this effect on species richness differed among genotypes (testing the interaction between genotype and sample type, F=3.66; df=4, 67; p=0.009), and marginally so for Shannon diversity (F=2.41; df=4, 67; p=0.057; Figure 4). Separate analyses on the *Daphnia* gut microbiomes and bacterioplankton showed no significant effect of culture type (mono- versus cocultures), species (*D. magna* vs *D. pulex*), genotypes or their interactions (Table SI2) on species richness and Shannon diversity.

**Figure 3:**
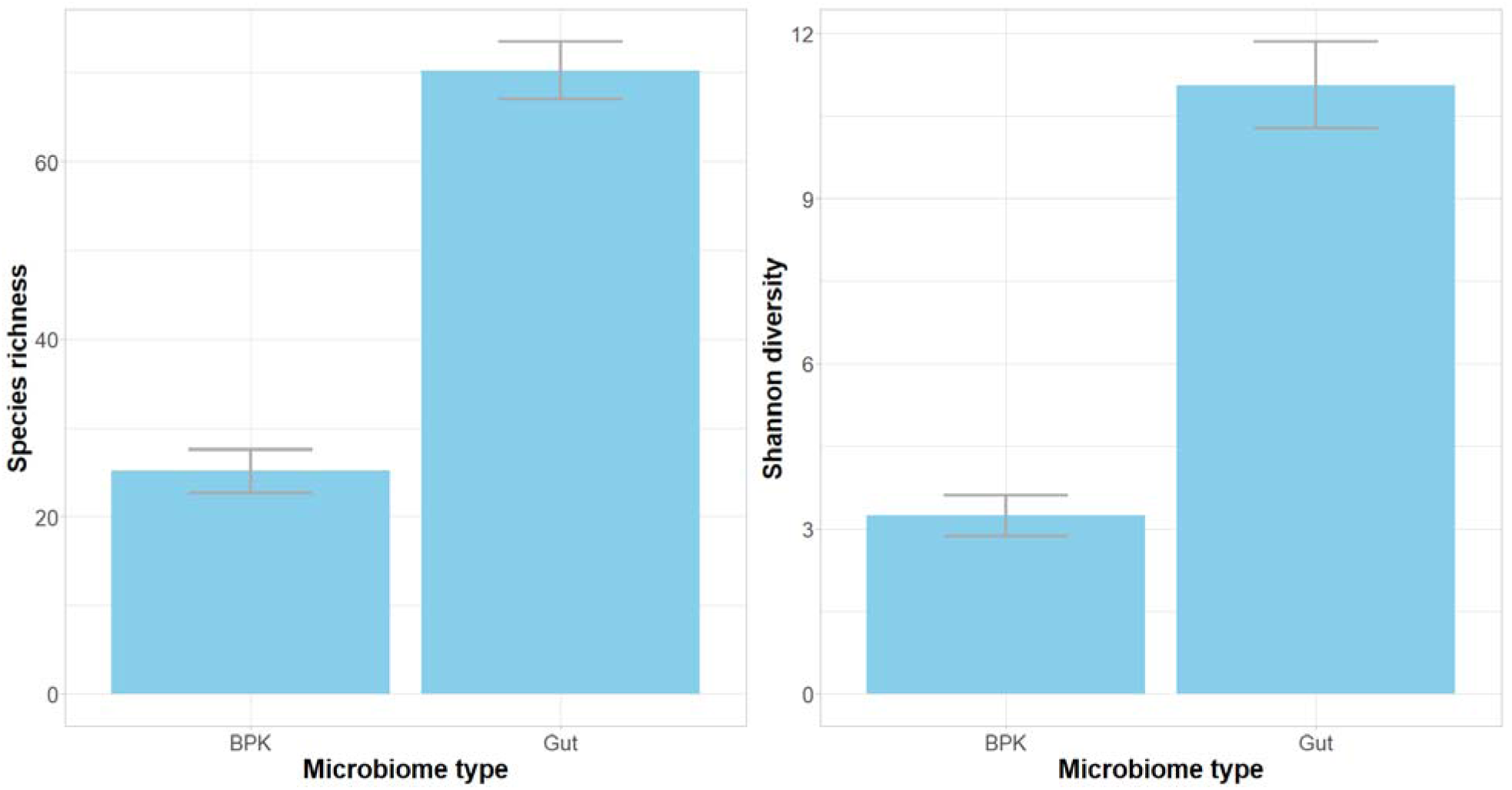
Species richness (left) and Shannon diversity (right) of the bacterioplankton (BPK) and gut microbiomes (GUT). Error bars represent one standard error of the mean.

**Figure 4:**
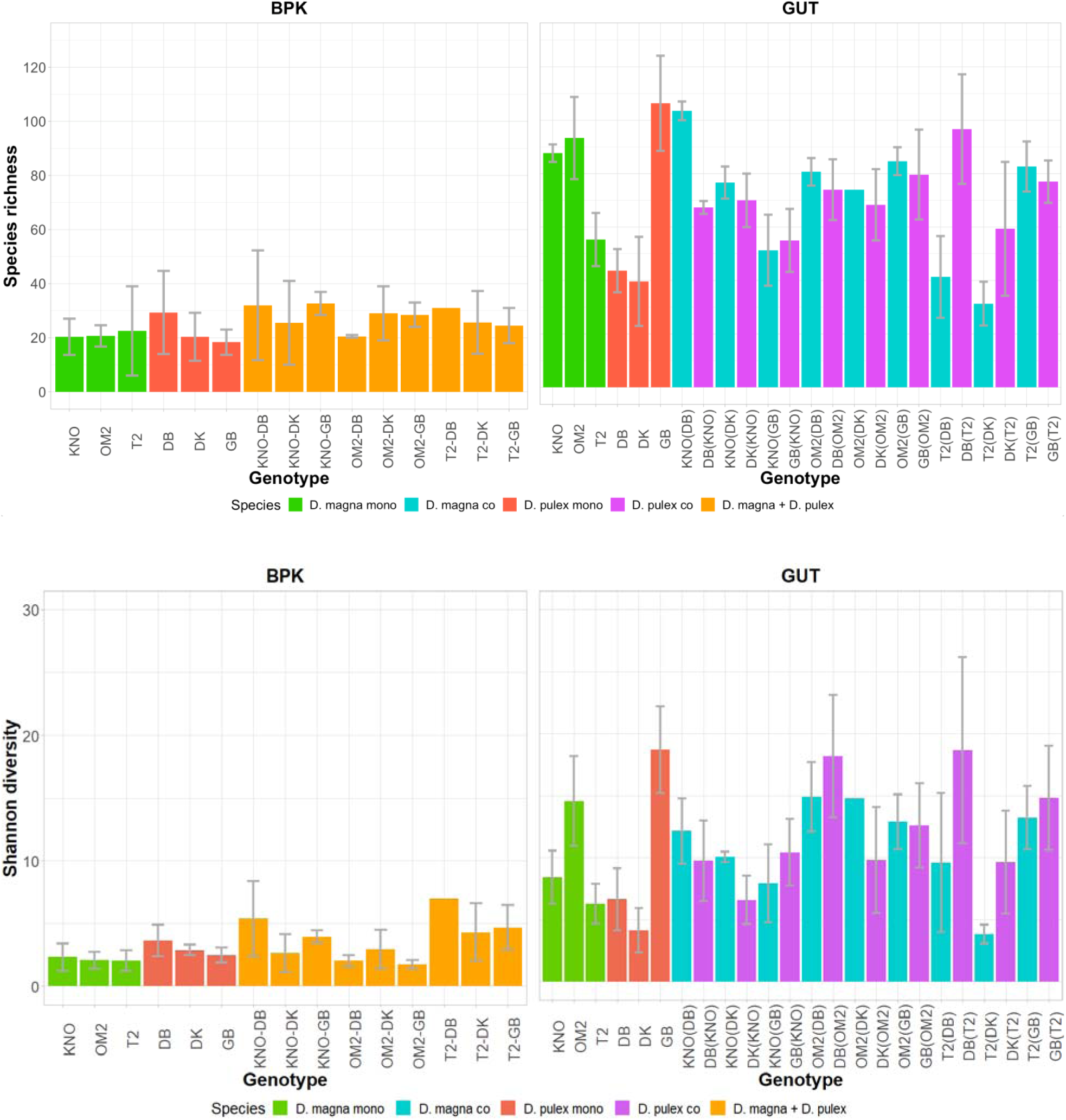
Species richness (upper row) and Shannon diversity (bottom row) of the bacterioplankton (BPK, left) and gut microbiomes (GUT, right) of the different combinations of *D. magna* and *D. pulex* genotypes. Error bars represent one standard error of the mean. For the gut microbiomes, the representation of the genotypes is as follows: KNO(DB) are gut microbiomes from the *D. magna* genotype KNO that was in cocultures with *D. pulex* genotype DB. DB(KNO) are gut microbiomes from the *D. pulex* genotype DB that were in cocultures with *D. magna* genotype KNO. For two treatments (BPK: T2-DB and GUT: OM2(DK)) the error bars are missing, this is because only one replicum was left after rarifying the data (some samples were removed because of the low number of reads).

### Beta diversity

When analyzing the full dataset (bacterioplankton and gut microbiomes), the Bray-Curtis dissimilarity between communities evinced a significant difference in bacterial composition between the bacterioplankton and the gut microbiomes (F=65.16, R^2^= 0.39, df=1, 12.77, p=0.001; Figure 5). All other factors (and their interactions) were not significantly associated with Bray-Curtis dissimilarities between bacterial communities. Separate analyses of Bray-Curtis dissimilarities on the bacterioplankton and gut microbiomes showed no significant differences between mono- and cocultures, between both host species (*D. magna* and *D. pulex*), between the genotypes of a given host species, or using the interactions between these factors.

**Figure 5:**
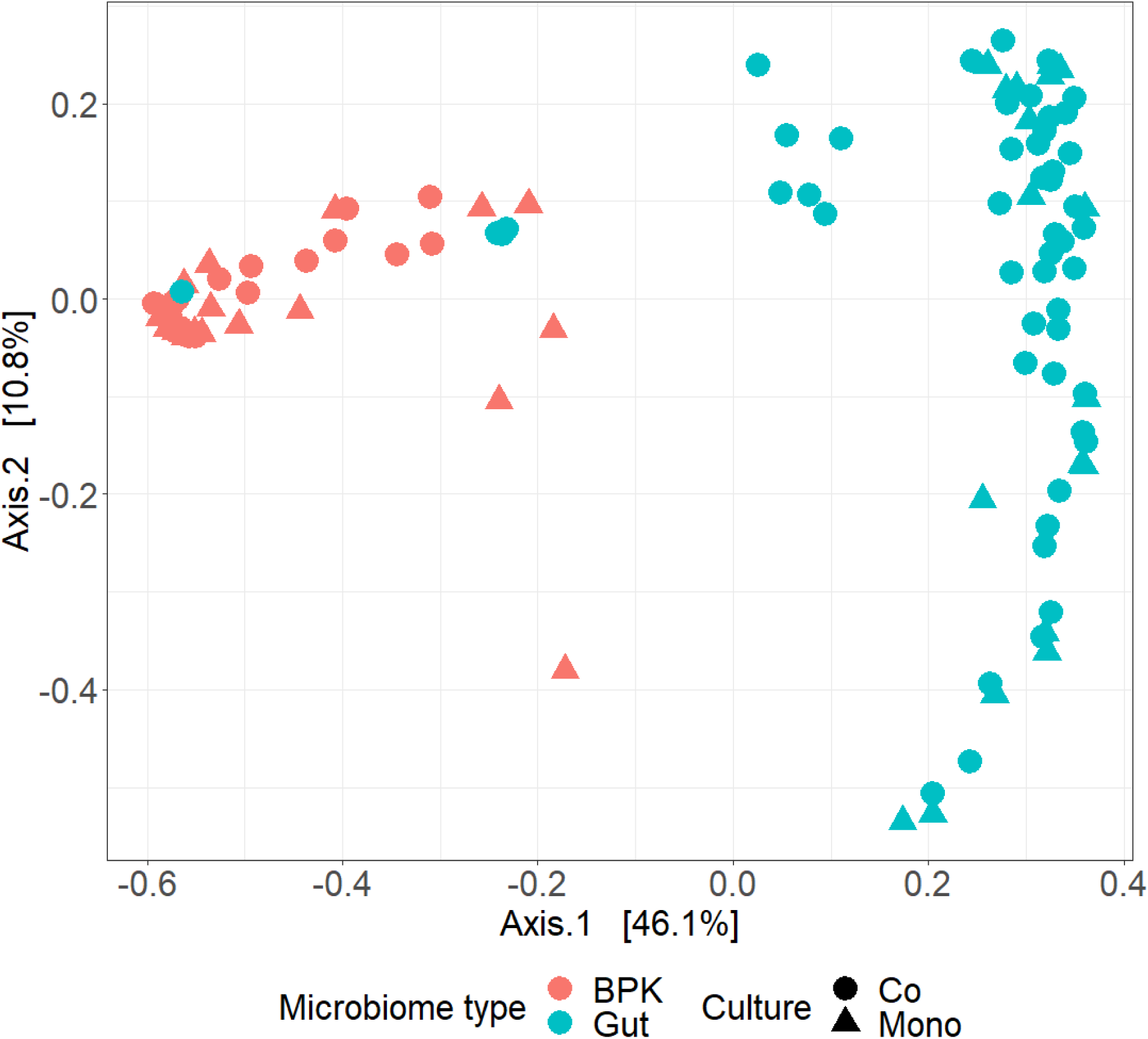
Ordination plot representing the beta diversity (Bray-Curtis distance) of bacterioplankton (BPK) and gut microbiomes (Gut). Cocultures are represented by dots and monocultures by triangles. The bacterioplankton samples are given in red and the gut microbiome samples in blue.

Dissimilarity indices (Bray-Curtis, weighted and unweighted Unifrac distances) were further analyzed to investigate the differences between the gut microbiomes of *D. magna* and *D. pulex* when they were cultured in mono- versus cocultures. Per genotype combination (all *D. magna* –*D. pulex* couples: KNO-DB, KNO-DK, KNO-GB, OM2-DB, OM2-DK, OM2-GB, T2-DB, T2-DK, T2-GB) the distance between the two species was calculated, both when they were in mono- and in cocultures. For all dissimilarity indices, there were significant differences (Table 2) between the two host species, among their genotypes, and also among genotype x host species combinations. In six of the nine *D. magna* genotype x *D. pulex* genotype combinations, the distances between the bacterial community compositions of the two species are higher when in monoculture than when in coculture (Figure 6 upper row, Table 2 and SI7). Furthermore, six of the nine combinations show that when in cocultures, *D. pulex* were more like *D. magna,* while *D. magna* in cocultures were not (or to a lesser extent) more like *D. pulex* in monocultures (Figure 6 middle row, Table 2 and SI7). Finally, six of the nine genotype combinations show that for the genotypes in cocultures, *D. pulex* microbiomes changed more than the ones from *D. magna* (Figure 6 lower row, Table 2 and SI7). Noteworthy, each time when a *D. magna* genotype is combined with the *D. pulex* genotype GB, the trend is in the opposite direction. One possible explanation for this deviation is the high species richness in the *D. pulex* genotype GB in mono- cultures (Figure 4).

**Figure 6:**
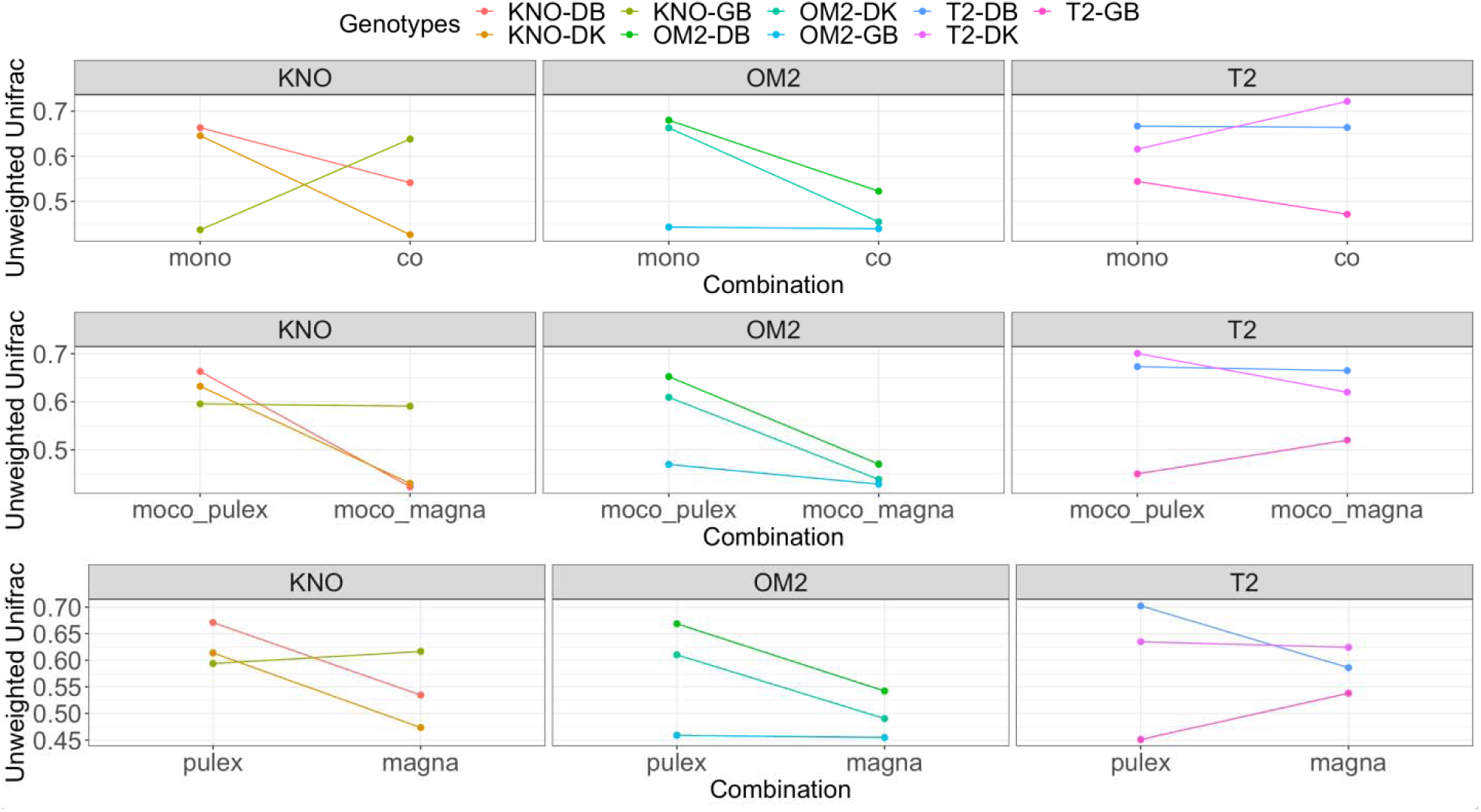
Overview of Unweighted Unifrac distances in microbiome community composition between *D. magna* and *D. pulex* for the different combinations of host species and culture type. Upper row: *D. magna* in monocultures versus *D. pulex* in monocultures (mono) and *D. magna* in cocultures versus *D. pulex* in cocultures (co). Middle row: Unweighted Unifrac distance between *D. magna* in cocultures versus *D. pulex* in monocultures (moco_pulex) and *D. magna* in monocultures versus *D. pulex* in cocultures (moco_magna). Lower row: Unweighted Unifrac distance between *D. pulex* in monocultures versus *D. pulex* in cocultures (pulex) and *D. magna* in monocultures versus *D. magna* in cocultures (magna). Distances between *D. magna* and *D. pulex* are represented per genotype. The plots are aligned in three columns that represent the three *D. magna* genotypes. The nine colors represent the different *D. magna-D. pulex* combinations.

**Table 2:**
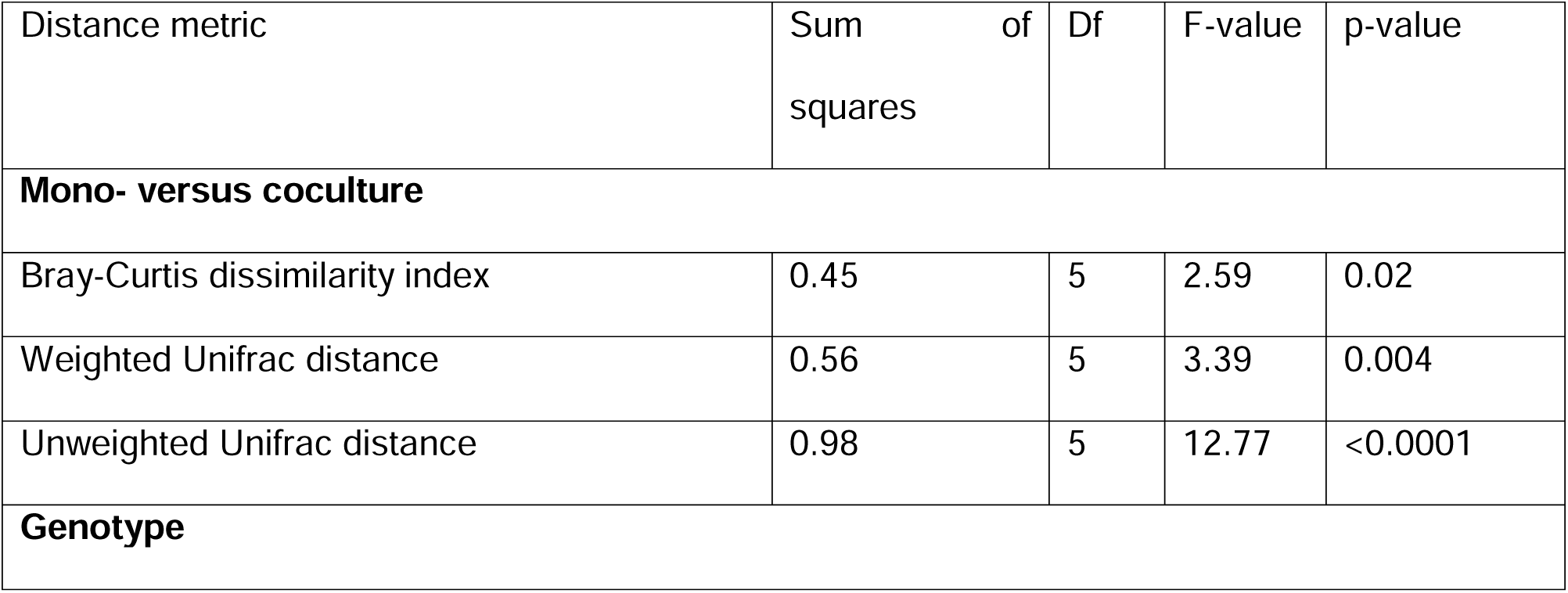
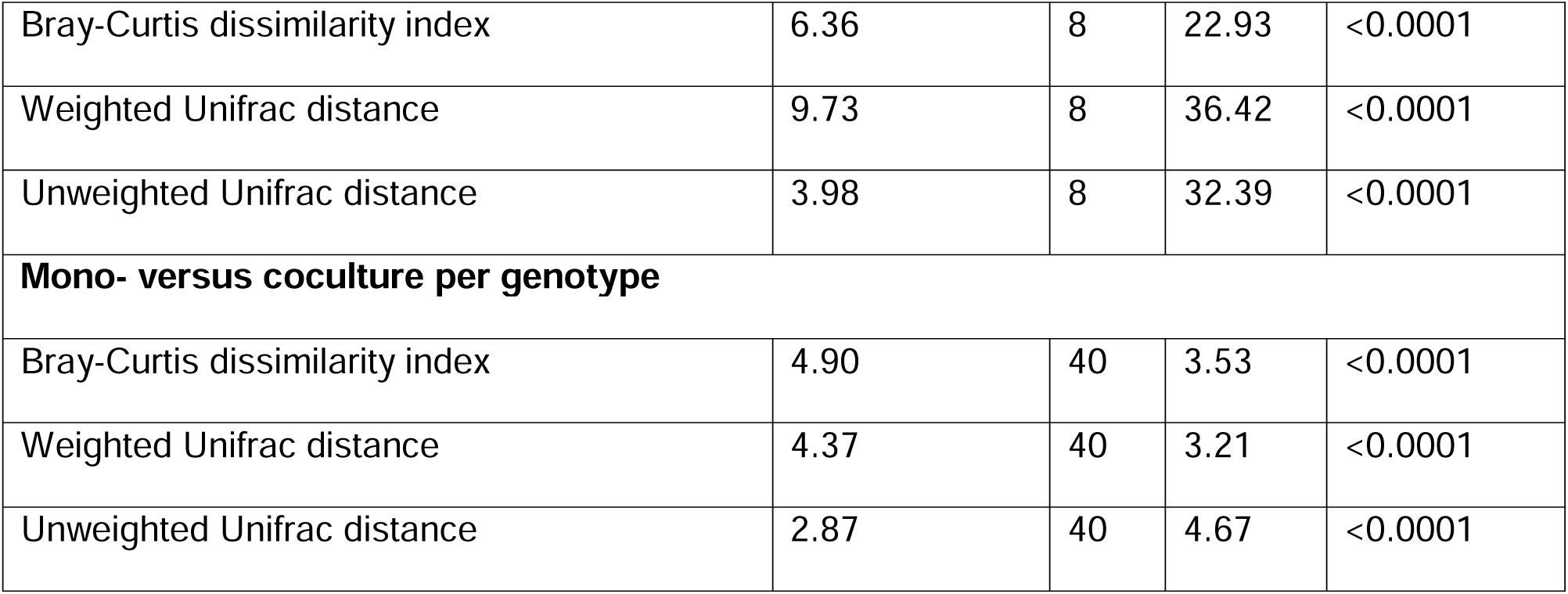
Results of the comparative analysis (*D. magna* versus *D. pulex*) of species composition between mono- and cocultures and between genotypes using a general linear model with an F-test (Type II Anova).

| Distance metric | Sum of squares | Df | F-value | p-value |
| --- | --- | --- | --- | --- |
| **Mono- versus coculture** |  |  |  |  |
| Bray-Curtis dissimilarity index | 0.45 | 5 | 2.59 | 0.02 |
| Weighted Unifrac distance | 0.56 | 5 | 3.39 | 0.004 |
| Unweighted Unifrac distance | 0.98 | 5 | 12.77 | <0.0001 |
| **Genotype** |  |  |  |  |
| Bray-Curtis dissimilarity index | 6.36 | 8 | 22.93 | <0.0001 |
| Weighted Unifrac distance | 9.73 | 8 | 36.42 | <0.0001 |
| Unweighted Unifrac distance | 3.98 | 8 | 32.39 | <0.0001 |
| **Mono- versus coculture per genotype** |  |  |  |  |
| Bray-Curtis dissimilarity index | 4.90 | 40 | 3.53 | <0.0001 |
| Weighted Unifrac distance | 4.37 | 40 | 3.21 | <0.0001 |
| Unweighted Unifrac distance | 2.87 | 40 | 4.67 | <0.0001 |

### Microbiome composition

The bacterioplankton community mainly consisted of Actinobacteria (53.38%) and Gammaproteobacteria (35.36%), while the more diverse gut microbiomes also contained Bacteroidia (28.95%) and Verrucomicrobiae (5.73%), next to Actinobacteria (21.50%) and Gammaproteobacteria (38%; Figure 7, Table SI3). The DESeq analysis showed that the relative abundance of eleven ASVs significantly differed between gut microbiomes and associated bacterioplankton samples (Table SI4). Investigation of differences between gut microbiome and bacterioplankton per genotype showed that for most genotypes the gut microbiome composition differed from the bacterioplankton (Table SI5). Investigation of the bacterial differences among bacterioplankton samples showed no ASVs that significantly differed in relative abundance between mono- and cocultures and between host species. There were, however, some ASVs that significantly differed in relative abundance when hosts were considered at the genotype level. More specifically, the bacterioplankton associated with *D. magna* genotypes in monocultures differing from that associated with the same genotypes in cocultures, while the bacterioplankton associated with *D. pulex* genotypes were similar in mono- and cocultures (Table SI6). Investigation of bacterial differences among the gut microbiomes showed no ASVs that significantly differed in relative abundance between the mono- and cocultures. There were, however, two ASVs that significantly differed in relative abundance between the species: the relative abundance of ASV7 (Verrucomicrobiaceae) was significantly higher in *D. pulex* monocultures than in *D. magna* monocultures, and the relative abundance of ASV73 (Rosermartima, Planctomycetaceae) was significantly higher in *D. pulex* cocultures compared with *D. magna* mono- and cocultures (list of ASVs with their respective name in Table SI8). At the genotype level, there were also some ASVs that significantly differed in relative abundance (Tables 3 and SI8). Table 3 shows a summary of the genotypes in mono- and cocultures where the relative abundance of some ASVs differed. In six of the nine pairwise combinations of *D. magna* with *D. pulex* genotypes, the gut microbiome of *D. pulex* in monocultures differed from the gut microbiome of either *D. magna* or *D. pulex* in cocultures. For the other three combinations, one showed no difference between *D. magna* and *D. pulex* in both mono- and cocultures and in the other two combinations *D. magna* in monocultures differed from *D. magna* or *D. pulex* in cocultures. Thus, in most cases (six out of nine) the gut microbiome of *D. pulex* genotypes (DB, DK and GB) in coculture changed in the direction of the gut microbiome of the *D. magna* genotypes (KNO, OM2 and T2).

**Figure 7:**
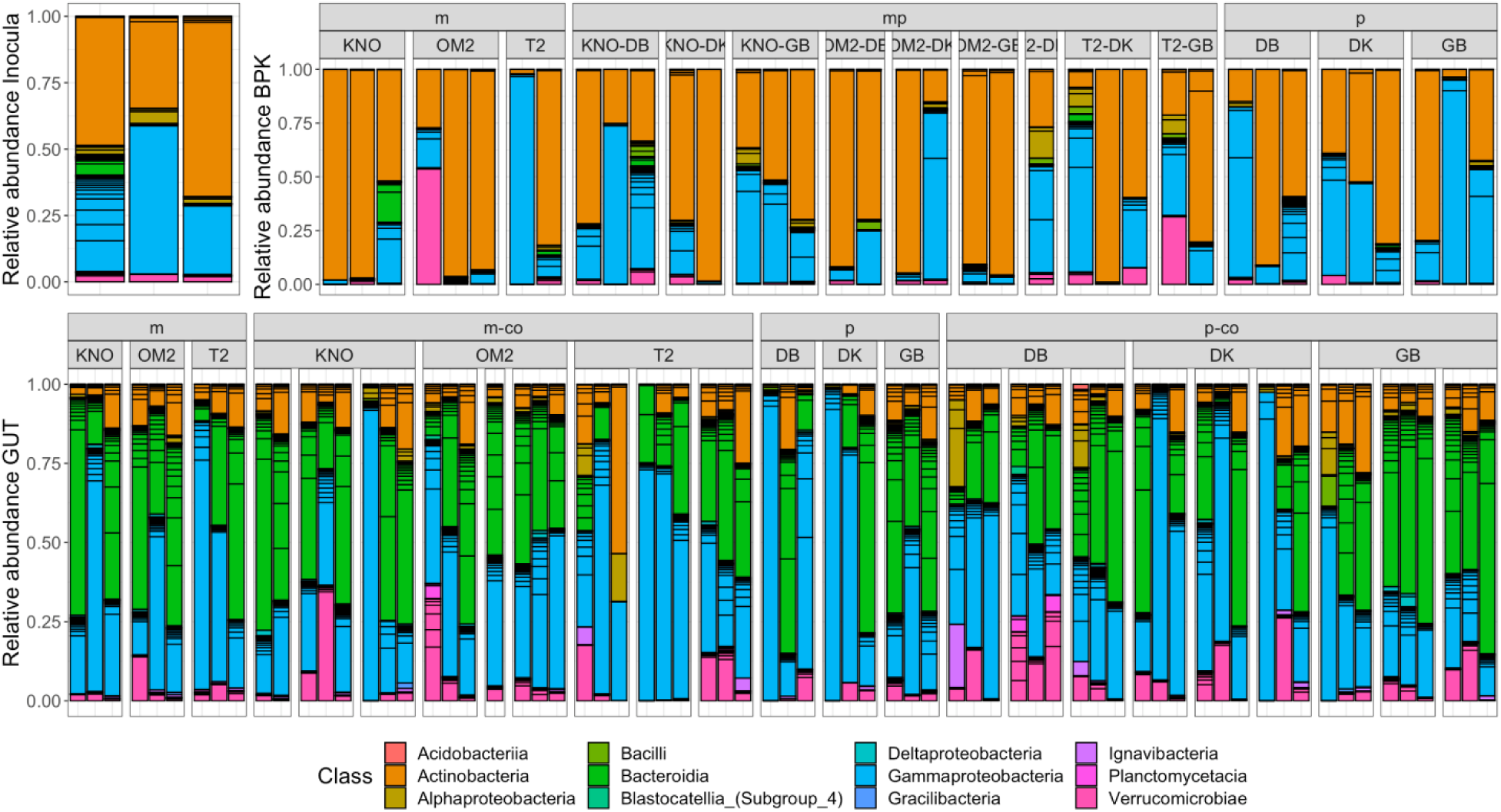
Relative abundance of the bacterial classes in the inocula (upper row, left), bacterioplankton (BPK, upper row, right) and gut microbiomes (GUT, lower row) of *D. magna* and *D. pulex* in mono- and cocultures. The samples are grouped per species (m, p, mp, m-co and p-co) and genotype (KNO, OM2, T2, DB, DK and GB). M = *D. magna* monocultures, p = *D. pulex* monocultures, mp = bacterioplankton of cocultures of *D. magna* and *D. pulex,* m-co = gut microbiome of *D. magna* cocultures, p-co = gut microbiome of *D. pulex* cocultures.

**Table 3:**
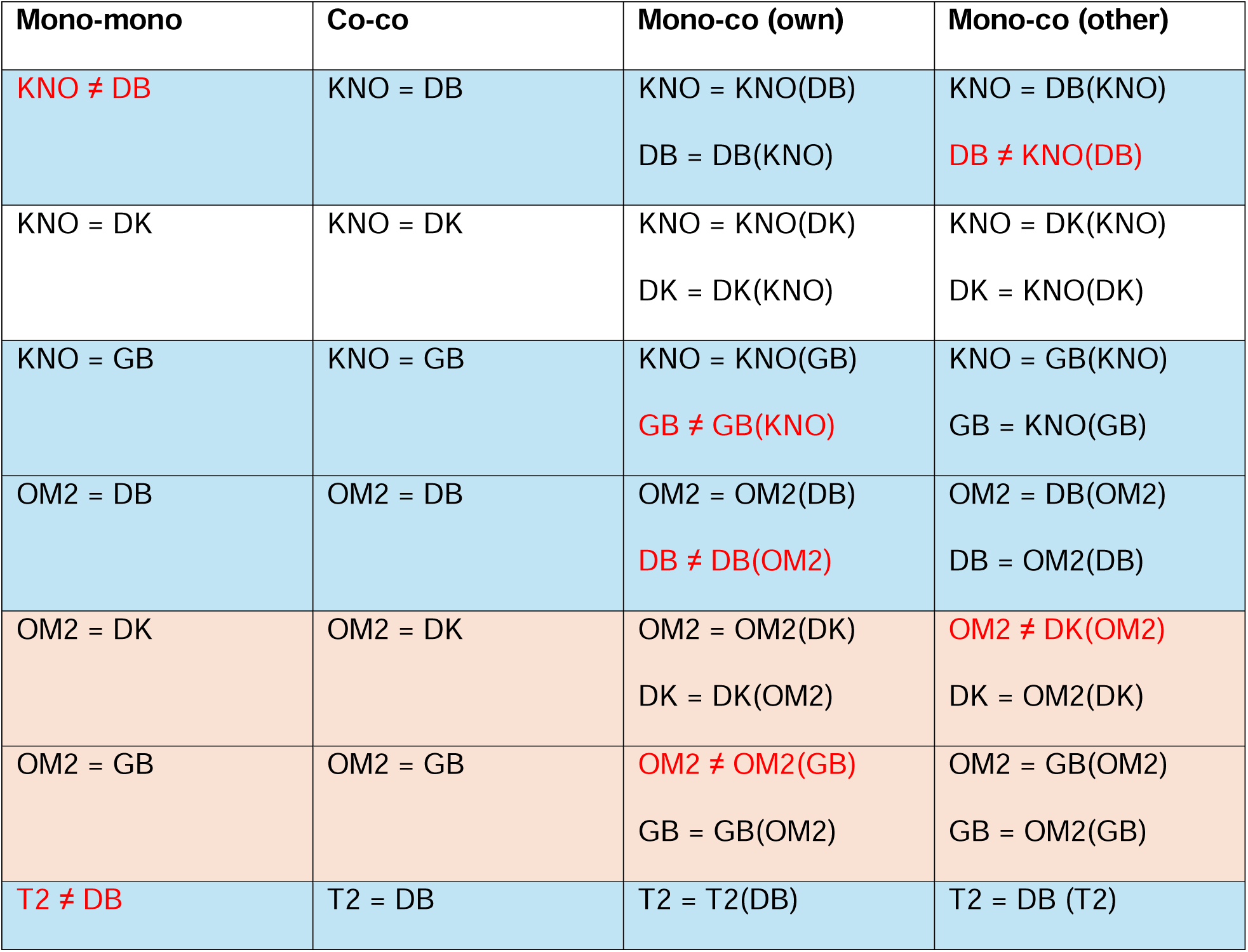
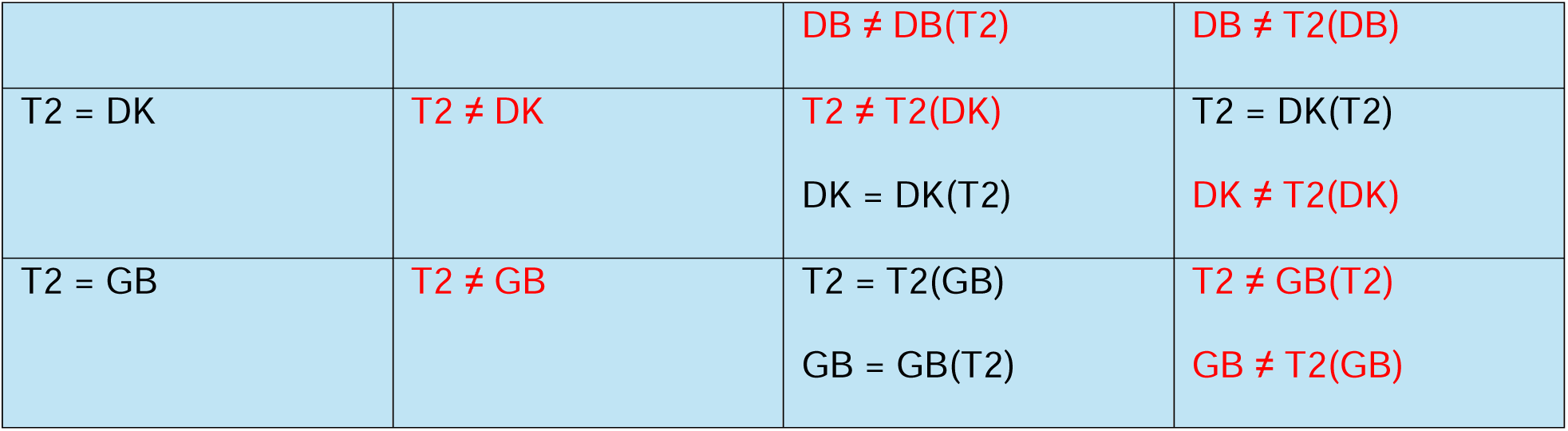
Summary of the pairwise comparison of the presence of significantly different relative abundances of ASVs in the gut microbiomes between genotypes (DESeq analysis). The table shows whether the relative abundance of at least one ASV was significantly different (≠) between the two genotypes compared (details in Table SI8). Relative abundances that differed significantly between two genotypes are in red. Relative abundances that are equal between samples are in black. Orange shaded cells refer to cases where *D. magna* in monocultures differed from *D. magna* or *D. pulex* in cocultures. Blue shaded cells refer to cases where *D. pulex* in monocultures differed from *D. magna* or *D. pulex* in cocultures. Representation of the genotypes is as follows: KNO(DB) are gut microbiomes from the *D. magna* genotype KNO that was in cocultures with *D. pulex* genotype DB. DB(KNO) are gut microbiomes from the *D. pulex* genotype DB that was in cocultures with *D. magna* genotype KNO.

The most abundant ASV in the bacterioplankton was ASV1: Mycobacterium (Table 4), with the highest relative abundance in *D. magna* monocultures (64.53%), followed by cocultures (55.35%), and *D. pulex* monocultures (46.48%). This ASV was also the most abundant ASV in the gut microbiomes (Table 5), with the highest relative abundance in *D. magna* in coculture (14.23%), followed by *D. pulex* in cocultures (14.12%), *D. magna* in monocultures (13.19%) and *D. pulex* in monocultures (7.91%).

**Table 4:** Relative abundance of the 10 most common ASVs in the bacterioplankton, grouped per host species, with m= *D. magna* in monocultures, p= *D. pulex* in monocultures, mp= cocultures.

| ASV | m | mp | p |
| --- | --- | --- | --- |
| ASV_1_Mycobacterium_sp. | 64.53 | 55.35 | 46.55 |
| ASV_4_Betaproteobacteriales | 9.21 | 12.22 | 16.62 |
| ASV_2_Ralstonia_sp. | 4.31 | 7.76 | 9.51 |
| ASV_10_Burkholderiaceae | 2.94 | 0.87 | 3.90 |
| ASV_8_Verrucomicrobiaceae | 5.13 | 1.54 | 0.67 |
| ASV_11_Mycobacterium_sp. | 0.87 | 1.13 | 1.32 |
| ASV_18_Hydrogenophaga_sp. | 0.57 | 0.81 | 1.42 |
| ASV_26_Betaproteobacteriales | 0.15 | 0.79 | 0.98 |
| ASV_61_Betaproteobacteriales | 0.62 | 0.56 | 0.65 |
| ASV_21_Mycobacterium_sp. | 0.63 | 0.45 | 0.56 |

**Table 5:** Relative abundance of the 10 most common ASVs in the gut microbiomes, grouped per species, with m= *D. magna* in monocultures, p= *D. pulex* in monocultures, m-co= *D. magna* in cocultures, and p-co= *D. pulex* in cocultures.

| ASV | m | m-co | p | p-co |
| --- | --- | --- | --- | --- |
| ASV_1_Mycobacterium_sp. | 13.19 | 14.23 | 7.91 | 14.13 |
| ASV_3_env.OPS_17 | 15.03 | 9.52 | 7.21 | 10.10 |
| ASV_5_Caenimonas_sp. | 7.44 | 6.36 | 7.09 | 8.38 |
| ASV_7_Burkholderiaceae | 3.36 | 3.14 | 13.41 | 1.90 |
| ASV_2_Ralstonia_sp. | 5.14 | 6.24 | 0.21 | 6.00 |
| ASV_8_Verrucomicrobiaceae | 2.47 | 3.07 | 1.58 | 3.91 |
| ASV_10_Burkholderiaceae | 2.53 | 3, | 2.01 | 2.22 |
| ASV_12_Flavobacterium_sp. | 1.33 | 2.22 | 3.57 | 2.27 |
| ASV_9_Kocuria_sp. | 0.03 | 0.02 | 7.46 | 0.04 |
| ASV_15_Burkholderiaceae | 2.06 | 1.67 | 2.66 | 0.88 |

The Venn diagram of the bacterioplankton (Figure 8) showed that cocultures had the highest percentage of unique ASVs (35.9%), followed by *D. pulex* monocultures (7.1%) and *D. magna* monocultures (4.9%). The Venn diagram of the gut microbiomes (Figure 9) showed that the gut microbiome of *D. pulex* in cocultures had the highest percentage of unique ASVs (8.2%), followed by *D. magna* in cocultures (3.9%), *D. magna* in monocultures (1.2%) and *D. pulex* in monocultures (1.2%). In addition, the two species shared more gut ASVs in cocultures (76.2%) than between their respective monocultures (46.8%).

**Figure 8:**
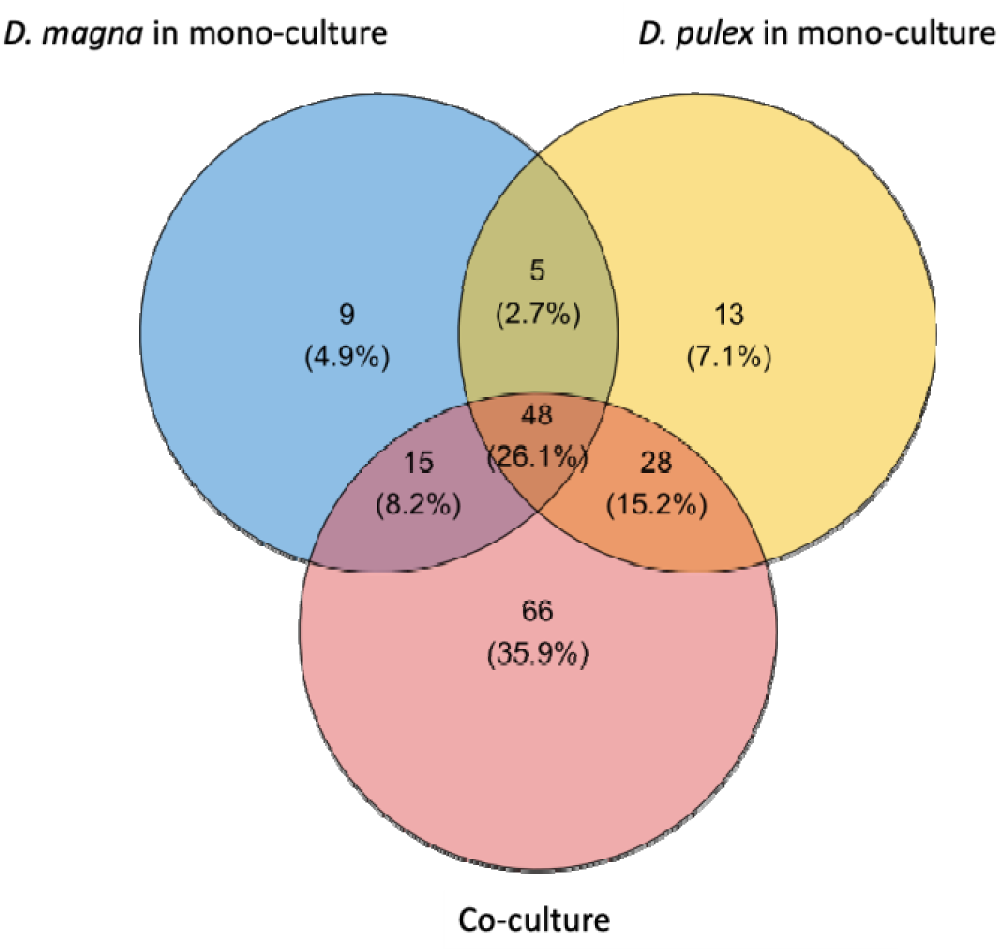
Venn diagram of the bacterioplankton, showing the unique and shared ASVs for *D. magna* in monocultures, *D. pulex* in monocultures and cocultures. The numbers show the number of ASVs and the relative abundance in percentage of the ASVs per subgroup.

**Figure 9:**
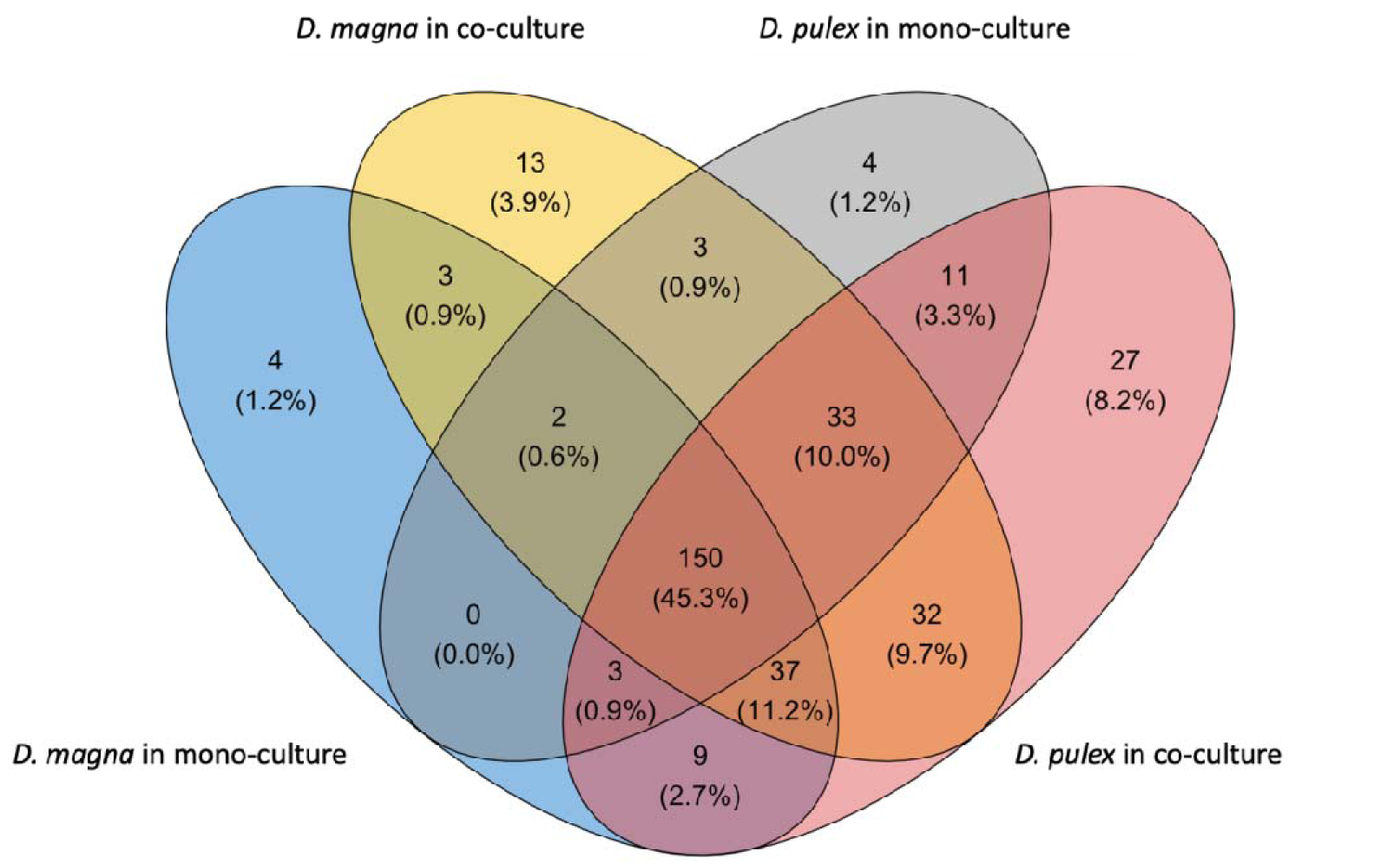
Venn diagram representing the unique and shared ASVs of the gut microbiomes, grouped per species and culture type (based on the rarified data). The numbers show the number of ASVs and the relative abundance in percentage of the ASVs per subgroup.

## Discussion

In this study, we experimentally investigated the effect of interspecific co-occurrence on the gut microbiome composition and diversity of two frequently co-occurring *Daphnia* species, *D. magna* and *D. pulex*. We expected that the gut microbiome would differ between *D. magna* and *D. pulex*, and that the larger filter-feeder *D. magna* would have a larger effect on the smaller filter-feeder *D. pulex* in cocultures compared to monocultures than *vice versa* (Figure 1). The working hypothesis was that *D. magna* would affect the gut microbiome of the weaker competitor *D.pulex*, most likely by affecting the bacterioplankton composition. The latter is possible through grazing of the bacterial population affecting the abundance of dominant strains, through enrichment of certain strains via defecation, or through facilitative effects in the *D. pulex* microbiome through early colonizing *D. magna* bacterial strains resulting in more similar microbiomes of both species in co- than in monocultures.

The diversity of the gut microbiomes was higher than that of the bacterioplankton. This was reflected in both the alpha (species richness and Shannon diversity) and the beta diversity. Previous studies showed that zooplankton (inclusive *Daphnia*) gut microbiomes differed from the bacterioplankton community (Macke et al., 2017, 2020, Callens et al., 2020, Cooper and Cressler, 2020, Samad et al., 2020). Results on difference in diversity between the bacterioplankton and the *Daphnia* gut microbiome are, however, not consistent. In experiments where germ-free *Daphnia* were inoculated with a microbial inoculum, as in this experiment, the gut microbiome community is often more diverse than the surrounding bacterioplankton (Macke et al., 2017; 2020 exp 2). In contrast, if *Daphnia* are not made germ-free before inoculation with a microbial inoculum, than diversity in the bacterioplankton is higher than in the *Daphnia* gut microbiome (Macke et al., 2020 exp 1; Coone et al., 2023). Most likely this effect depends on the colonizing bacteria differing in their establishment success and growth inside the *Daphnia* gut, which can be driven by abiotic factors, the host immune system or competitive interactions with other bacteria (Callens et al., 2020; Bulteel et al., 2021; Cooper and Cressler, 2020; Eckert et al., 2021). Interestingly in this respect is that sterilized *Daphnia* individuals inoculated with sympatric gut microbiomes (suggesting more adaptive interactions between the host and the microbiome) have a higher gut microbiome diversity than *Daphnia* individuals inoculated with allopatric gut microbiomes when exposed to toxic cyanobacteria (Houwenhuyse et al., 2021). Also Pielou evenness (Figure SI8) was higher in the *Daphnia* gut microbiomes than in the bacterioplankton. This means that bacterial strains were more evenly distributed in the *Daphnia* gut microbiomes than in the bacterioplankton. In addition, the relative abundance tables of the 10 most common ASVs in the bacterioplankton (Table 4) and gut microbiomes (Table 5) showed that some ASVs had a higher relative abundance in the *Daphnia* gut than in the bacterioplankton (i.e., ASV3: env.OPS_17, ASV5: *Caenimonas* sp., ASV7: Burkholderiaceae sp., ASV12: *Flavobacterium* sp., ASV9: *Kocuria* sp., and ASV15: Burkholderiaceae).

Differences in the gut microbiome composition among species can occur when, for example, the species differ in their niche, in that way they encounter a different pool of microbial species (Adiar et al., 2020, Eckert et al. 2021, Schols et al. 2023). In terms of alpha and beta diversity, we did not observe strong differences between the two species *D. magna* and *D. pulex*. Overall, there was no strong evidence for the existence of a species-specific gut microbiome. This agrees with Qi et al. (2009) comparing different laboratory strains of multiple *Daphnia* species. Also, Eckert et al. (2021) did not find a strong signal for zooplankton host species when they examined species specificity of microbiomes in a field study. They suggested that filter-feeding zooplankton can continuously recruit bacteria from the surrounding water, which is advantageous as freshwater environments are highly fluctuating and pelagic bacteria at a specific moment are already adapted to such conditions. There is probably redundancy of beneficial bacterial functions in the environment, which makes it difficult to detect strict host-microbiome associations (Eckert et al., 2021). Important external factors that can influence the gut microbiome of *Daphnia* are (i) physical environmental factors such as temperature (Sullam et al., 2018, Frankel-Bircker et al., 2019; Samad et al. 2020), (ii) diet (Macke at al., 2017), and (iii) the environmental pool of bacteria (Callens et al., 2020; Macke et al., 2020). The two *Daphnia* species used in this study, *D. magna* and *D. pulex*, have a very similar diet (Bengtsson, 1986), habitat occupation (DeMott and Pape, 2005), and anatomy (Kuster and von Elert, 2013), which might explain why only subtle differences in gut microbiome composition and diversity were present between these two *Daphnia* species.

But even being subtle, we hypothesized that the gut microbiome composition of one species would depend on the presence of the other species, in other words, that the gut microbiome composition of *D. magna* and *D. pulex* in cocultures would differ from those found in their monocultures. For the majority of the genotype combinations: (i) the distance in microbiome composition (DESeq analysis) between *D. pulex* and *D. magna* was higher in monocultures than in cocultures; (ii) the distance between *D. pulex* in monocultures and *D. magna* in cocultures was higher than the distance between *D. magna* in monocultures and *D. pulex* in cocultures; (iii) the distance between *D. pulex* in monocultures and *D. pulex* in cocultures was higher than the distance between *D. magna* in monocultures and *D. manga* in cocultures; and (iv) *D. pulex* in monocultures differed more in microbiome composition from either *D. magna* or *D. pulex* in cocultures than *D. magna* in monocultures from either *D. magna* or *D. pulex* in cocultures. This tentatively suggests that the microbiome of *D. pulex* (the weaker filter feeder) becomes more like *D. magna* (the stronger filter feeder) when cultured together. *D. pulex* in cocultures had the highest number of unique ASVs. This suggests that the weaker competitor *D. pulex* took up more unique bacteria in its gut in the presence of a stronger, filter-feeding competitor. This is in accordance with the fact that stronger grazing (by the stronger competitor) may select for rare, grazing resistant strains (Jürgens et al., 1999), but further testing is needed to elucidate this. Another possible explanation for these differences in microbiome composition between *D. magna* and *D. pulex* in mono- and cocultures is the difference in biomass. As *D. magna* is larger bodied than *D. pulex*, *D. magna* has more room to host (*D. magna* specific?) bacteria in their gut, which in turn may lead to a seemingly stronger influence of *D. magna* on *D. pulex*, than vice versa.

Finally, we investigated whether differences in the gut microbiomes between mono- and cocultures were mediated by the bacterioplankton. The subtle differences between mono- and cocultures observed in this experiment, suggest some level of host microbiome sharing between the two species via the surrounding environment (bacterioplankton) and potentially also microbiota-microbiota interactions between the present and incoming bacteria (as suggested in Zélé et al., 2018; Chong et al., 2019; Decaestecker, Van de Moortel et al., 2024). In our data, the most dominant ASV from the bacterioplankton (ASV1: *Mycobacterium* sp.) had a higher relative abundance in *D. magna* than in *D. pulex*, when comparing monocultures and cocultures separately (*D. magna:* monocultures: 13.19% versus cocultures: 14.23%, *D. pulex*: monocultures: 7.91% versus cocultures: 14.12%). This, in addition with the observation that *D. pulex* in cocultures had the highest number of unique ASVs, tentatively suggests that the larger filter-feeder, *D. magna,* filtered out the most dominant/abundant bacteria, which were then taken up by *D. pulex*. This result is in accordance with Callens et al. (2018) who showed that high doses of antibiotics reduced the abundance of the most abundant/dominant strains in *D. magna*, resulting in a more diverse community as other taxa could increase in the absence of the most dominant strains. This also suggests that aquatic host associated microbiomes are open systems, with hosts and environments reciprocally affecting each other’s microbial composition (Macke et al., 2020; Callens et al, 2020). Members of the host’s microbiome can spread to the environment and serve as a microbial source for other hosts (Castro-Sanguino & Sánchez, 2012; Sweet, 2014; Chandler et al., 2011; Mistry et al., 2017).

We conclude that in general, the composition of the gut microbiome did not differ between *Daphnia magna* and *Daphnia pulex* when all genotypes are considered. We did, however, observe subtle differences with respect to the gut microbiome composition analyses at the genotype level. For most of the genotype combinations (six out of nine), the microbiome of *D. pulex* changed more when cultured in cocultures than in monocultures in comparison with *D. magna*. Our results also tentatively suggest that the stronger filter-feeder *D. magna* filtered out the most abundant/dominant ASVs from the environment, leaving the bacterioplankton with unique, grazing resistant strains that were then taken up by the weaker filter-feeder *D. pulex*. But further testing is needed to prove this.

